# Asymmetric molecular architecture of the human *γ*-tubulin ring complex

**DOI:** 10.1101/820142

**Authors:** Michal Wieczorek, Linas Urnavicius, Shih-Chieh Ti, Kelly R. Molloy, Brian T. Chait, Tarun M. Kapoor

## Abstract

The *γ*-tubulin ring complex (*γ*-TuRC) is an essential regulator of centrosomal and acentrosomal microtubule formation ^1–4^. Metazoan *γ*-TuRCs isolate as ∼2 MDa complexes containing the conserved proteins *γ*-tubulin, GCP2 and GCP3, as well as the expanded subunits GCP4, GCP5, and GCP6 ^3,5,6^. However, in current structural models, *γ*-TuRCs assemble solely from subcomplexes of *γ*-tubulin, GCP2 and GCP3 ^7^. The role of the metazoan-specific subunits in *γ*-TuRC assembly and architecture are not currently known, due to a lack of high resolution structural data for the native complex. Here, we present a cryo-EM structure of the native human *γ*-TuRC at 3.8Å resolution. Our reconstruction reveals an asymmetric, single helical-turn and cone-shaped structure built from at least 34 polypeptides. Pseudo-atomic models indicate that GCP4, GCP5 and GCP6 form distinct Y-shaped assemblies that structurally mimic GCP2/GCP3 subcomplexes and are distal to the *γ*-TuRC “seam”. Evolutionary expansion in metazoan-specific subunits diversifies the *γ*-TuRC by introducing large (>100,000 Å^2^) surfaces that could interact with different regulatory factors. We also identify an unanticipated structural bridge that includes an actin-like protein and spans the *γ*-TuRC lumen. Despite its asymmetric composition and architecture, the human *γ*-TuRC arranges *γ*-tubulins into a helical geometry poised to nucleate microtubules. The observed compositional complexity of the *γ*-TuRC could self-regulate its assembly into a cone-shaped structure to control microtubule formation across diverse contexts, e.g. within biological condensates ^8^ or alongside existing filaments ^9^.

## Main text

The *<u>γ</u>*<u>-tu</u>bulin <u>r</u>ing <u>c</u>omplex (*γ*-TuRC) is a large macromolecular assembly that is essential for proper microtubule network formation in eukaryotes ^1–4^. Our understanding of *γ*-TuRC structure comes mainly from studies of heterotetramers of *<u>γ</u>*-tubulin <u>c</u>omplex <u>p</u>rotein <u>2</u>, or GCP2, GCP3, and *γ*-tubulin, also termed the *<u>γ</u>*<u>-tu</u>bulin <u>s</u>mall <u>c</u>omplex (*γ*-TuSC) ^5^. In *S. cerevisiae*, the *γ*-TuSC is a Y-shaped complex whose base is comprised of laterally associated N-terminal domains of GCP2 and GCP3 ^10^. Notably, recombinant *S. cerevisiae γ*-TuSCs have a propensity to self-assemble into helical oligomers of varying length *in vitro* ^7^. However, negative stain electron tomography and metal shadowing studies have indicated that *D. melanogaster γ*-TuRCs, which are biochemically similar to the human complex, assemble into finite, lock washer-shaped structures ^11^. In addition to GCP2 and GCP3, human *γ*-TuRCs also contain the metazoan-specific proteins GCP4, GCP5, and GCP6 ^6,12,13^, but structural models for any of the core subunits in the context of the native complex are not known. Further, despite ∼30 years of research since the discovery of *γ*-tubulin ^14^, it is unclear whether the human *γ*-TuRC assembles from a helical oligomer of *γ*-TuSCs or even adopts a lock-washer shaped structure.

To examine the architecture of human *γ*-TuRCs, we first sought to purify the native complex. Several previously reported purification strategies were explored ^3,6^. We found that isolation of the native *γ*-TuRC from HeLa S3 cytoplasmic extracts via its affinity for the CM1 domain of CDK5RAP2, also termed the *<u>γ</u>*<u>-Tu</u>RC-mediated <u>N</u>ucleation <u>A</u>ctivator (*γ*-TuNA) ^15^, reliably yielded complex containing core components whose proportions matched previous qualitative estimates ^6^, as judged by SDS-PAGE and iBAQ analysis of mass spectrometry data (Figure S1; see Methods for purification details) ^16^.

We determined the structure of the native human *γ*-TuRC using cryo-electron microscopy (cryo-EM) to an overall resolution of 3.8 Å (Figure 1A and Figure S2; see Methods for details). The *γ*-TuRC structure revealed a conical shape ∼30 nm in diameter and ∼20 nm in height (Figure 1A). 14 similarly-sized (∼4 nm-wide) globular densities lined the open face of the cone, each supported by a stalk-like structure. Both the globular domains and the lower parts of the stalk-like structures made substantial lateral contacts with one another, while leaving several narrow “gaps” in the conical wall of the complex (Figure 1A). Additionally, one of the stalk-like densities appeared to extend down and plug the opening found at the base of the *γ*-TuRC (the “plug” in Figure 1A). The combination of two neighboring stalk-like densities along with their two associated globular domains closely resembled previous descriptions of the Y-shaped *γ*-TuSC ^10^ (Figure 1A). The human *γ*-TuRC cone shape is therefore made of 14 individual subunits that could form 7 *γ*-TuSC-like subcomplexes (Figure 1A-C).

**Figure 1.**
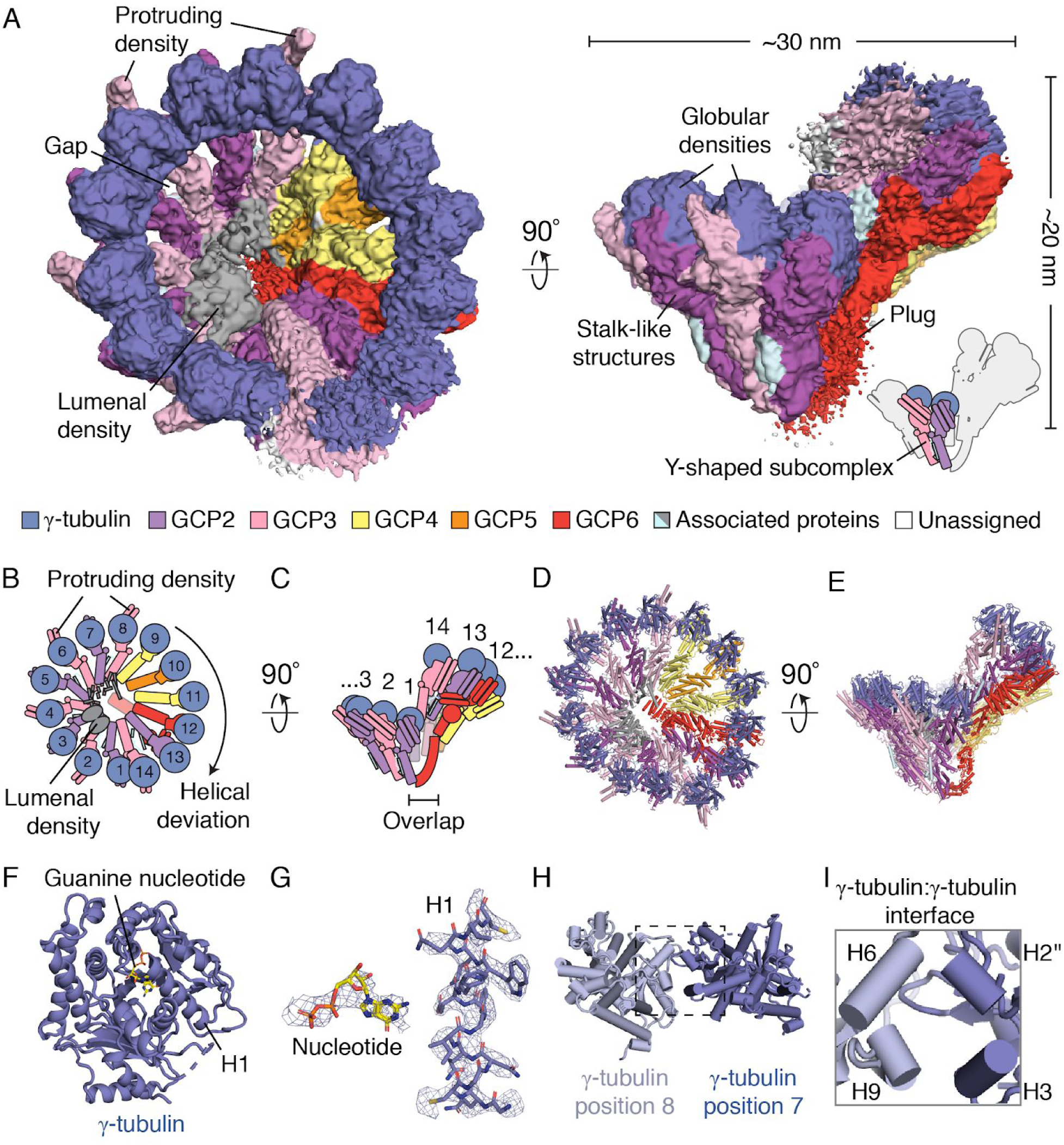
Cryo-EM reconstruction of the native human *γ*-TuRC. A) Two views of the overall *γ*-TuRC density map (surface representation). Structural features and dimensions of the *γ*-TuRC are indicated. Schematic of the *γ*-TuRC highlighting a Y-shaped subcomplex (indicated) is shown in the bottom right. B) and C) Schematics of the *γ*-TuRC highlighting proposed subunit numbering, features of asymmetry (both compositional and structural), and the “overlap” region. D) and E) Two views of a *de novo* molecular model for the *γ*-TuRC (cartoon representation). F) Refined *γ*-tubulin model (cartoon representation) with guanine nucleotide (stick representation) and helix H1 indicated. G) Examples of *γ*-tubulin density quality. Refined models for GDP (left, stick representation) and helix H1 of *γ*-tubulin (right, stick representation) shown in the corresponding *γ*-tubulin density (blue mesh). H) Model of the *γ*-tubulin:*γ*-tubulin interface at positions 8 (light blue cartoon representation) and 7 (blue cartoon representation). I) Inset from H) showing interacting helices (labeled cylinders). Except for H) and I), *γ*-TuRC subunits are colored according to the legend in A).

We numbered the individual stalk-like subunits, beginning at the base of the *γ*-TuRC cone and finishing at subunit 14, which partially overlapped with the first subunit (the *γ*-TuRC “overlap”) (Figure 1B-C). This organization denoted apparent helical symmetry in the complex. However, the *γ*-TuRC exhibited at least three features that revealed structural asymmetry (Figure 1A). First, an externally-protruding density appeared only at positions 2, 4, 6, and 8 of the complex (“protruding densities”; Figure 1A), suggesting that a set of four similar *γ*-TuSC-like subcomplexes, each with one protruding density, occupied these positions. Second, the remaining three *γ*-TuSC-like subunits lacking the protruding densities and approaching the overlap region of the cone (positions 9 to 14) deviated from a perfect helical geometry, conferring ellipticity to the *γ*-TuRC cone shape (the “helical deviation”; Figure 1, A-B). Third, a substantial (∼10 x 2 x 4 nm) density was situated in the lumen of the cone structure and contacted the stalk-like densities from positions 2-10 (the “lumenal density”; Figure 1A). These results suggest that, in contrast to previous models ^17^, the human *γ*-TuRC adopts an asymmetric conical structure comprised of heterogeneous *γ*-TuSC-like subunits.

The *γ*-TuRC density map, which was generated from ∼460,000 particles (Figure S3), displayed a broad range of local resolution (3.0 Å - 12 Å; Figure S2C), revealing a degree of structural “breathing” in this large complex. To improve the local resolution across the *γ*-TuRC and gain detailed insight into its molecular features, we processed different parts of the complex separately in a “divide and conquer” strategy that employed the use of particle subtraction ^18^ (Figure S3; see Methods for processing details). In brief, signal in raw particles not corresponding to desired segmentation regions in the *γ*-TuRC was subtracted. Signal-subtracted particles were then subjected to focused 3D classification and refinement. The resulting locally refined, segmented maps had more homogeneous resolution (FSC_0.146_ ranged from 3.4 to 4.5 Å for 6 of the 7 maps, and 6.5 Å for the one map corresponding to the “overlap” region; Figure S3). These maps revealed clearly-defined side chain densities across the majority of complex and were used to generate a near-complete molecular model of the core *γ*-TuRC structure (Figure 1D-E).

The structure of *γ*-tubulin in the context of the *γ*-TuRC is not currently known. Previous high-resolution structural models for *γ*-tubulin come from crystallographic studies of the monomeric human protein ^19,20^. We rigid-body fitted a crystallographic model for human *γ*-tubulin into the 14 globular densities at the open face of the cone described above (Figure 1A; PDB ID: 3CB2 ^20^). After refinement, 13 of these densities clearly accommodated *γ*-tubulin side chains, as well as a guanine nucleotide (Figure 1F-G). The conformations of “hallmark” structural features (*α*-helices H6-H7 and the intermediate β-sheet) of our refined *γ*-tubulin model closely resembled the original crystallographic model (RMSD = 1.0 Å; Figure S4A-C) ^19,20^. These data suggest that, under our purification conditions (see Methods), and in contrast with a recent study of a recombinant yeast *γ*-TuSC ^21^, assembly into the native human *γ*-TuRC does not induce significant structural changes in *γ*-tubulin.

Our results also provided details about the *γ*-tubulin:*γ*-tubulin lateral interface, which includes interactions between helix H2’’ and H3 of one *γ*-tubulin with helix H6 and H9 of the subsequent *γ*-tubulin (Figure 1H & I). Surprisingly, we observed two repeating arrangements of these interfaces; in one, the neighboring *γ*-tubulins were ∼9 Å apart (between positions 1-2, 3-4, 5-6, 7-8, 9-10, and 11-12), and in the other, neighboring *γ*-tubulins were farther (∼15 Å) apart (between positions 2-3, 4-5, 6-7, 8-9, 10-11, and 12-13) (Figure S4D-E). We note that this alternating variation in the inter-*γ*-tubulin interfaces is consistent with different spacings within and between *γ*-TuSC-like subcomplexes in the *γ*-TuRC.

Next, we turned our attention to the stalk-like densities that complete the Y-shaped *γ*-TuRC subcomplexes. The stalk-like densities had a characteristic overall organization comprising two bundles each built from ∼7 consecutive *α*-helices. The lower (N-domain) bundle adopted an elongated (∼8 x ∼2 nm) fold while the upper (C-domain) bundle, which contacted *γ*-tubulin, was more compact (∼5 nm x ∼4 nm) and displayed a prominent (>2 nm) two-helix hairpin on the outer face of the cone (Figure 2A-C). The general topology of the stalk-like structures resembled a previously described crystallographic model for GCP4, the only metazoan-specific core subunit of the *γ*-TuRC with available structural data. GCP4 is organized into two *α*-helical bundles containing the conserved *<u>γ</u>*-tubulin <u>r</u>ing complex interacting <u>p</u>rotein motif <u>1</u> (GRIP1, corresponding to the N-domain) and the GRIP2 (C-domain) motifs ^22^. Notably, GCP4 has also been proposed to form part of a *γ*-TuSC-like subcomplex at the *γ*-TuRC “overlap” ^17,23,24^. Surprisingly, however, the crystallographic model for GCP4 could not be rigid-body fitted and refined into the *γ*-TuRC density map at the “overlap” region (i.e., positions 1 or 14; also referred to as the *γ*-TuRC “seam” ^25^), but instead fit into the densities at positions 9 and 11 (Figure 2C, D, H & I; ∼83% GCP4 sequence coverage). The resolution of our density map enabled unambiguous side chain identification that confirmed the identity of GCP4 at these positions of the *γ*-TuRC (Figure 2D & I). Further, we aligned our refined GCP4 model to the monomeric, crystallographic GCP4 model via their N-domains ^22^, which revealed that within the *γ*-TuRC the GCP4 C-domain is rotated by ∼10° towards the preceding subunit (Figure S4F), and helix 21 is rotated by ∼90° towards the GCP4-associated *γ*-tubulin (Figure S4G). Together, these results indicate that the *γ*-TuRC contains two copies of GCP4, both of which are located distal to the “overlap” region of the complex.

**Figure 2.**
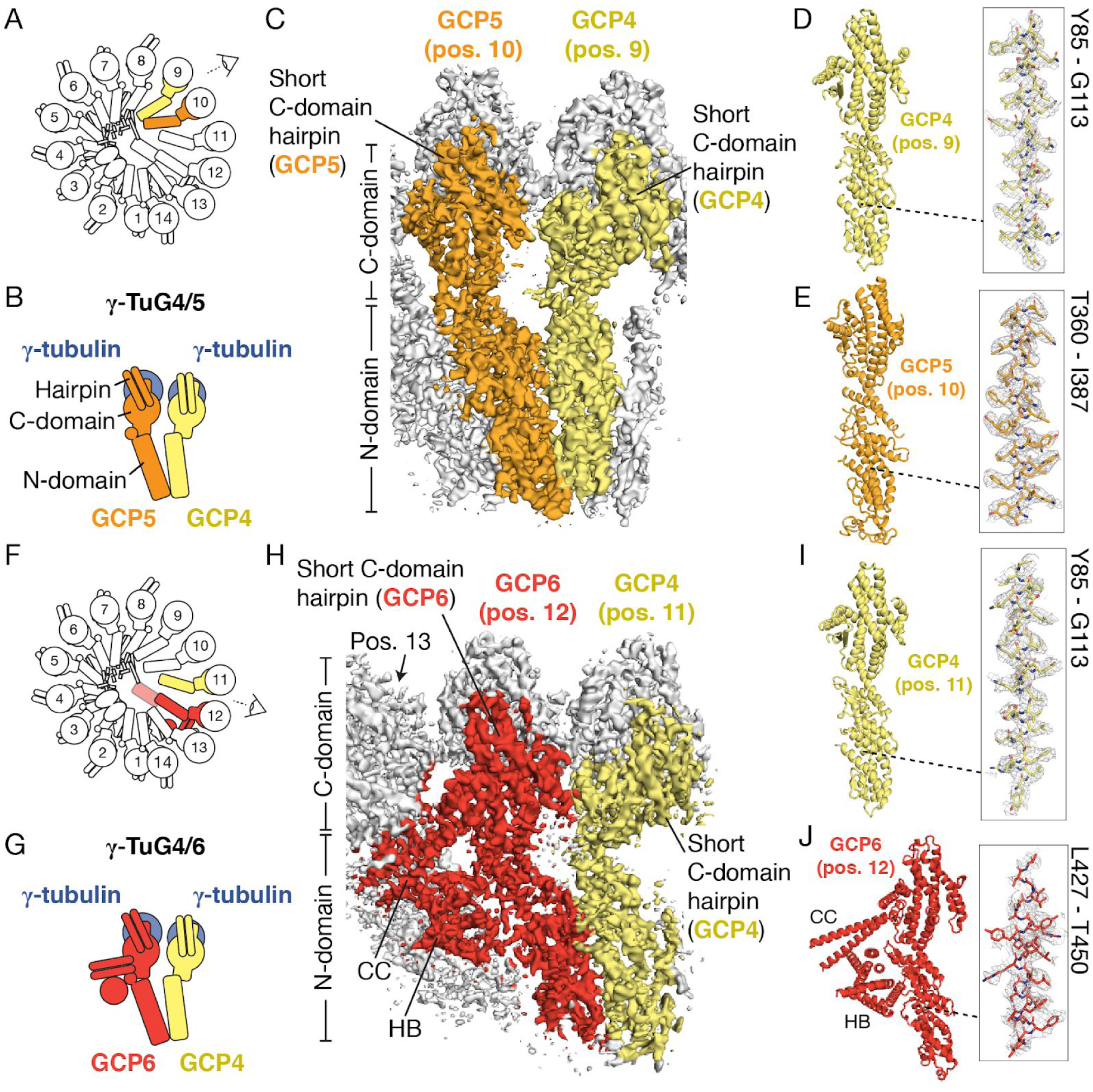
Asymmetric positioning of GCP4, GCP5 and GCP6 in the *γ*-TuRC. A) Schematic of the *γ*-TuRC establishing the viewing angle and highlighting the locations of GCP4 (yellow) and GCP5 (orange). B) Schematic of the *γ*-tubulin/GCP4/GCP5 heterotetramer that forms a Y-shaped *γ*-TuSC-like assembly, *γ*-TuG4/5. C) The density map of *γ*-TuRC positions 9-11 used to model N- and C-domains and short C-domain hairpins (indicated) of GCP4 (yellow surface) and GCP5 (orange surface), viewed from the angle indicated in A). D) and E) Molecular models (left) of GCP4 and GCP5. Insets (right) show the fit of a region of the models (indicated) in the density maps (mesh). F) Schematic of the *γ*-TuRC highlighting the locations of GCP4 (yellow) & GCP6 (red). G) Schematic of a *γ*-tubulin/GCP4/GCP6 heterotetramer that forms the *γ*-TuSC-like assembly, *γ*-TuG4/6. H) Density at positions 11-13 used to model N- and C-domains and short C-domain hairpins (indicated) of the second GCP4 (yellow surface) and GCP6 (red surface), viewed from the angle indicated in F). GCP6-associated coiled-coil (CC) and helical bundle (HB) densities interacting with position 13 are indicated. I) Molecular model of GCP4 at position 11 (left). Inset (right) shows the fit of GCP4 residues Y85-G113 (indicated; stick representation) in the density map (mesh). J) Molecular model of GCP6 at position 12 (left), including poly-alanine models of the CC and HB densities (indicated). Inset (right) shows the fit of GCP6 residues L427-T450 (indicated; stick representation) in the density map (mesh).

The remaining metazoan-specific subunits, GCP5 and GCP6, are also proposed to be situated at the *γ*-TuRC “overlap” region ^17^, but their structure, position and stoichiometry within the complex are not known. Further, in addition to the conserved GRIP1 (N-domain) and GRIP2 (C-domain) motifs that define the GCP family ^26^, GCP5 and GCP6 also feature large sequence inserts (described below; Figure S5A and Figure S6A) ^6^. We first focused on building a model for GCP5, as it has previously been shown to biochemically interact with GCP4 ^23^. Consistent with this report, we found that a *de novo* molecular model for human GCP5 N- and C-domains could be built into the stalk-like density at position 10 of the *γ*-TuRC (Figure 2E and Figure S5A). As above, analysis of amino acid side chain registries in *α*-helical densities enabled unambiguous assignment of GCP5 to this position of the complex (Figure 2E). Notably, however, the GCP5 model did not fit into the stalk-like densities at the *γ*-TuRC “overlap” (positions 1, 2, 13, or 14), nor did this protein fit into any other stalk-like density in the complex. Our model also revealed that, consistent with conservation of the GRIP1/GRIP2 motifs, the core GCP5 model is similar in overall topology to GCP4, except for slight conformational variability in the C-domain (Figure S4H). Following the naming convention of the *γ*-TuSC ^5^, we propose that GCP4 at position 9 and GCP5 at position 10, along with their associated *γ*-tubulins, constitute a Y-shaped *γ*-TuSC-like subcomplex (*<u>γ</u>*<u>-tu</u>bulin subcomplex containing <u>G</u>CP<u>4</u> and GCP<u>5</u>, or “*γ*-TuG4/5”) (Figure 2B). Together, our results reveal the stoichiometry (1 per complex) and structural model of the N- and C-domains of GCP5 within the *γ*-TuRC, and demonstrate that GCP5 is located distal to the *γ*-TuRC “overlap”.

The GCP5 (position 10) density had two additional features that differed from other *γ*-TuRC stalk-like densities. First, 3D classification of the overall *γ*-TuRC density map revealed two subclasses that differed in the conformation of GCP5. The first subclass corresponded to the characteristic “elongated” stalk-like conformation that was used to build the GCP5 model described above (Figure S5C). In the second subclass, the GCP5 density adopted a “compacted” conformation within the complex, in which the GCP5-associated *γ*-tubulin was displaced ∼5-10 Å towards the base of the *γ*-TuRC (Figure S5D). Further, in the “compacted” GCP5 subclass, the proximal *γ*-TuSC-like subcomplexes occupying positions 11-14 were also displaced slightly away from the center of the *γ*-TuRC cone (Figure S5E-F). Second, we observed unassigned density below the GCP5 N-domain that interacts with the lumenal density (see below) as well as the neighboring GCP4 at position 9 (Figure S5B). This “N-terminal density” connects to our GCP5 model at a.a. 214-240 and likely corresponds in part to the ∼210 N-terminal amino acids currently not modeled in the GCP5 N-domain (Figure S5A).

We next focused on building a molecular model for GCP6, the remaining and largest of the metazoan-specific subunits ^6^. Again, in contrast with current models, which place this subunit at the *γ*-TuRC “overlap”, we found that a *de novo* model for the GCP6 N-domain could only be built and refined into the stalk-like density at position 12 (Figure 2H & J, and Figure S6A). The registry of model side chains matched the high-resolution densities in the N-domain region (Figure 2J), which, combined with the generally low sequence identity across the GCP N-domains ^6^, further validated the identity of GCP6 at this position. For the C-domain, unambiguous registry assignments using side chains was difficult due to less resolved density in this region of the *γ*-TuRC. Nonetheless, a secondary structure model could be built for the GCP6 C-domain based on secondary structure predictions and partial homology modeling using GCP4 as a template (see Methods for details) (Figure 2J and Figure S6A). Given the organizational similarity to the *γ*-TuG4/5 described above, we propose that the second GCP4 at position 11 and GCP6 at position 12, along with their associated *γ*-tubulins, constitute a distinct Y-shaped *γ*-TuSC-like subcomplex, the “*γ*-TuG4/6” (Figure 2G). Together, these results provide a molecular model for the N- and C-domains of GCP6 and indicate that, as with GCP4 and GCP5, GCP6 is positioned away from the *γ*-TuRC “overlap” region.

We next addressed the less-conserved regions of GCP6 (position 12). First, we noticed that the “plug” density in the overall map connects to the lower part of the GCP6 N-domain (Figure 1A and Figure S6B). A homology model for the remaining N-terminal residues of GCP6, which were not accounted for in the N-domain model, was generated based on a sequence-profile alignment ^27^ to a human spliceosomal protein (SF3b155; PDB ID: 5IFE ^28^; hereafter, the GCP6 “plug”; Figure S6C). Remarkably, this model matched the size and shape of the “plug” density (Figure S6D). Second, we observed a coiled coil density (CC) that stemmed from between the N- and C-domains of GCP6 at position 12 and contacted a small (∼2 nm wide) *α*-helical bundle (HB). Both the CC and HB in turn interacted with the neighbouring stalk density at position 13 (Figure 2H & J, and Figure S8B). Though currently unassigned, the large (∼800 residue) GCP6 insertion sequence located between the N- and C-domain models is predicted to form a coiled coil and may in part account for the CC and HB features (Figure 2J and Figure S6A).

Having accounted for the metazoan-specific core *γ*-TuRC subunits, we next sought to build models for the evolutionarily-conserved proteins GCP2 and GCP3, which are known to form *γ*-TuSCs in other species ^5,29^, but whose structures in the context of the *γ*-TuRC are not known. We found that near complete (∼72% full sequence coverage) *de novo* models for human GCP2 could be built into the stalk-like densities at positions 1, 3, 5 and 7 (Figure 3A, C & D, and Figure S7A). Similarly, GCP3 could be built into positions 2, 4, 6, and 8 (Figure 3A, C, & E, and Figure S7B). Side chain registries in these high-resolution densities confirmed the alternating positioning of these two proteins in the *γ*-TuRC (Figure 3D-E). Following historical convention ^5^, we refer to the Y-shaped arrangement of GCP2, GCP3 and their corresponding *γ*-tubulin molecules as the human “*γ*-TuSC” (Figure 3B & G).

**Figure 3.**
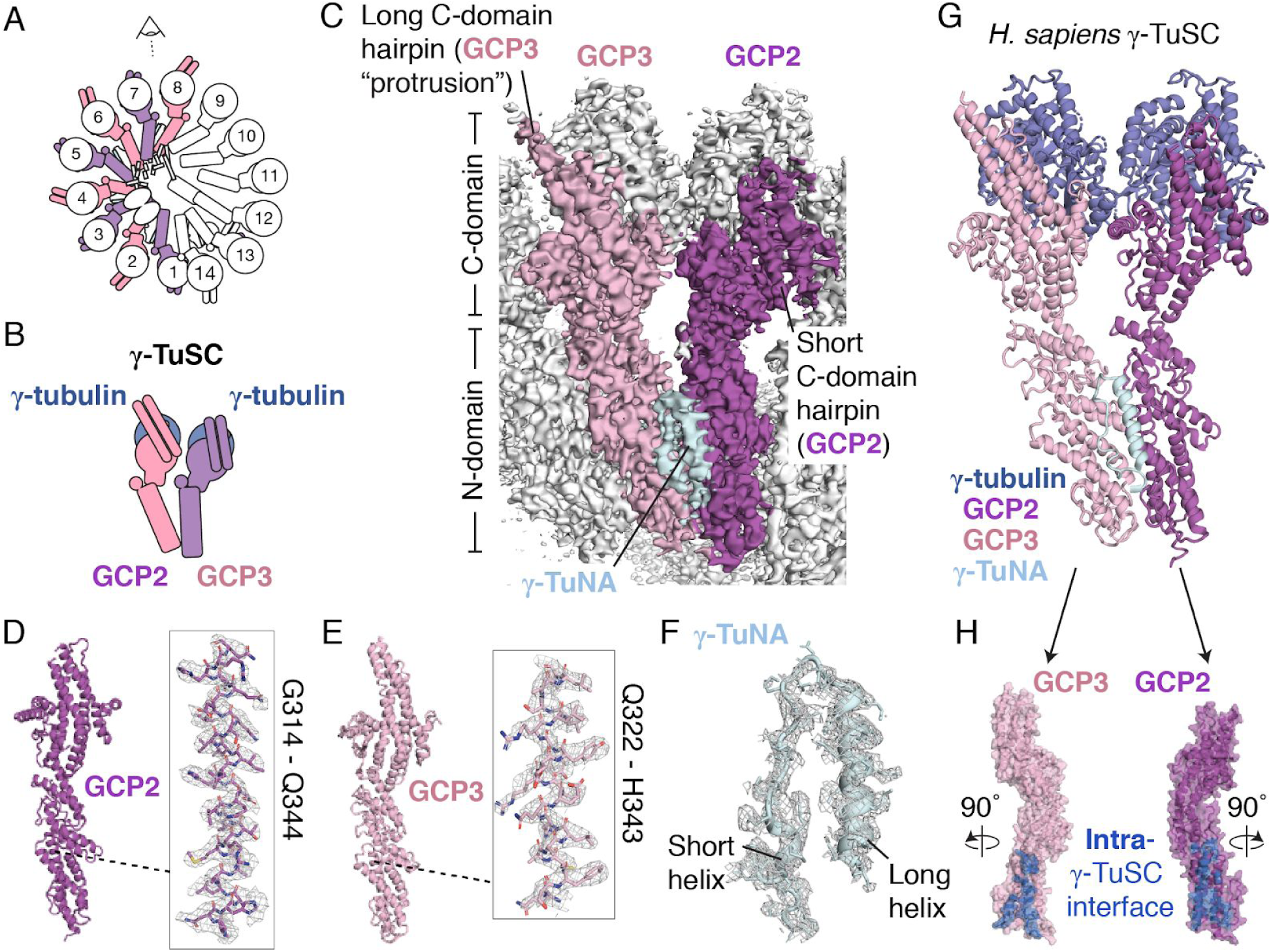
Structures and arrangement of human GCP2, GCP3 and the *γ*-TuNA. A) Schematic of the *γ*-TuRC highlighting locations of GCP2 (purple) and GCP3 (pink). B) Schematic of a *γ*-tubulin/GCP2/GCP3 heterotetramer that forms the human *γ*-TuSC. C) Overall density map used to model GCP2 (position 7, purple surface), GCP3 (position 8, pink surface), and the *γ*-TuNA (light blue surface) viewed from the angle indicated in A). N- and C-domains, long (GCP3) and short (GCP2) C-domain hairpins, and the *γ*-TuNA density, are indicated. D) Molecular model of GCP2 (left). Inset shows the fit of GCP2 residues G314-Q344 (stick representation) in the density map (mesh). E) Molecular model of GCP3 (left). Inset shows the fit of GCP3 residues Q322-H343 (stick representation) in the density map (mesh). F) Molecular model of the *γ*-TuNA (overlayed cartoon and stick representations) shown in the density map (mesh). Secondary structure features are indicated. G) Molecular model of the human *γ*-TuSC comprising GCP2, GCP3, and two copies of *γ*-tubulin. The *γ*-TuNA is also shown to be consistent with the density map in C). H) Rotated GCP2 (purple surface) and GCP3 (pink surface) models highlighting the intra-*γ*-TuSC interface (blue surface).

The GCP2 and GCP3 models were largely similar to one another, with the exception of the C-domain hairpin, which was ∼10 Å longer in GCP3 vs. GCP3 (Figure S7C), and which forms the basis of the repeating “protruding density” described in Figure 1. Further, our high-resolution models revealed details of a ∼3,000 Å^2^ interface between GCP2 and GCP3 (intra-*γ*-TuSC; Figure 3H), and between GCP3 and GCP2 (inter-*γ*-TuSC; Figure S7F). In addition, on the lumenal side of the *γ*-TuRC, a long (>50 Å) *α*-helix was found to form additional contacts with the N-domains of GCP2 and GCP3 at positions 3-6 (Figure S7G), which may stabilize interactions between multiple *γ*-TuSCs.

Unlike the *γ*-TuG4/5 and *γ*-TuG4/6, each human *γ*-TuSC is also associated with a small (∼10-20 Å) *α*-helical density found between the N-domains of GCP2 and GCP3 (Figure 3C). The location of this density is equivalent to the binding site for the CM1 domain-containing fragment of *S. cerevisiae* Spc110 (Spc110^1-220^) ^30^, which is crucial for the *in vitro* assembly of the budding yeast *γ*-TuSC ^7,10^. Guided by well-defined side chains in these *α*-helical densities, a *de novo* model for the CM1 domain of CDK5RAP2, also called the *γ*-TuNA ^15^, used in this study to purify *γ*-TuRCs, could be built into this region of each *γ*-TuSC (Figure 3F). The *γ*-TuNA model comprises a 45 residue helix-loop-helix motif that interacts with the N-domains of GCP2 and GCP3 near the intra-*γ*-TuSC interface (Figure S7D-E). Together, our results provide near-atomic structural models for the human *γ*-TuSC and the *γ*-TuNA, and indicate that four Y-shaped *γ*-TuSCs comprise the stalk-like densities at positions 1-8 of the human *γ*-TuRC.

We next sought to characterize the remaining subunits found at the *γ*-TuRC “overlap”. We found that position 13 of the *γ*-TuRC corresponds to an additional GCP2 based on the fit of side chains from our GCP2 model into the C-domain density (Figure 4A and Figure S8B-C), as well as the presence of a small *α*-helical density between the N-domains of positions 13 and 14 (Figure S8B), into which a rigid-body fit of our *γ*-TuNA model was possible. Next, we reasoned that the less-resolved stalk-like density at the subsequent position 14 likely corresponds to GCP3 based on a rigid body fit of our GCP3 model and the observation that GCP2 and GCP3 exist in pairs as part of the *γ*-TuSC ^5,31^ (Figure 4A and Figure S8E). Notably, the *γ*-TuSC and *γ*-TuSC-like subcomplexes at positions 1-12 are organized into a continuous lattice-like arrangement mediated by the lateral associations between GCP N-domains (Figure 1A). In contrast, the *γ*-TuSC at positions 13-14 is in a displaced configuration relative to the neighboring stalk-like subunits in the complex, in which the GCP2-associated *γ*-tubulin acts as a hingepoint for a ∼6° rotation of this *γ*-TuSC towards the conical axis (Figure 4, A and B). The rotation allows this *γ*-TuSC to avoid steric clashes that would otherwise occur in part between the N-domain of GCP3 (position 14) and the bi-lobed structure in the lumenal density (described below). Additionally, we observed a compact (∼2 x 2 x 3 nm), unassigned density that stemmed from the C-domain of GCP3 and was situated at the “overlap” region, just above the *γ*-tubulin at position 1 (Figure 4A-B, and Figure S8F). Together, our results indicate that, in contrast to previous models, the *γ*-TuRC “overlap” region is composed of two *γ*-TuSCs (one at positions 1-2, and another at positions 13-14).

**Figure 4.**
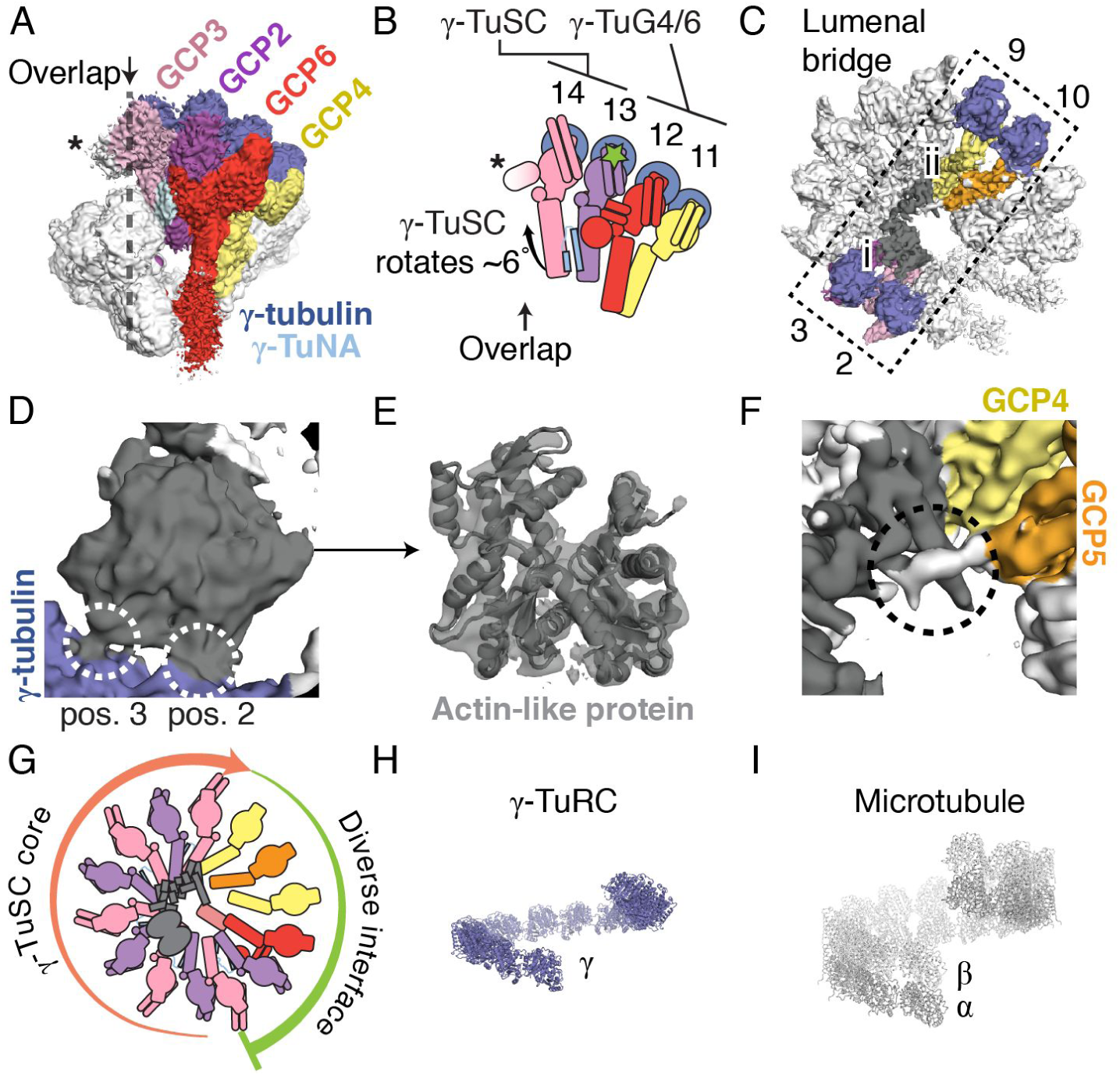
Asymmetric molecular architecture of the *γ*-TuRC. A) Side view of the *γ*-TuRC cryo-EM density map highlighting subunits leading up to the *γ*-TuRC “overlap” (indicated). A density at position 14 (asterisk) extending into the overlap from an arrangement of GCP2 & GCP3 (the “terminal *γ*-TuSC”) is indicated. B) Schematic of the *γ*-TuSC at the overlap adjacent to *γ*-TuG4/6 indicating its ∼6° rotation relative to a hinge point in the *γ*-tubulin at position 13 (green star). The GCP6 “plug” is omitted for clarity. C) Top view of the overall *γ*-TuRC density map highlighting the lumenal bridge (dashed rectangle, spanning from positions 2 and 3 to positions 9 and 10) and its two sub-domains (i: bi-lobed density domain; ii: *α*-helical bundle domain). D) Zoomed-in view of the interactions (white dashed circles) between the bi-lobed density ((i) in C); grey surface) and *γ*-tubulins at positions 2 and 3 (blue surface). E) A rigid-body fit of *β*-actin (grey cartoon representation; PDB ID 2HF3; ^32^) in the bi-lobed density (transparent grey surface). F) Zoomed-in view of the *α*-helical bundle domain ((ii) in C); grey surface) that interacts (dashed circle) with density (white surface) stemming from GCP5 (orange surface). G) Schematic of *γ*-TuRC organization. The *γ*-TuRC consists of two highly asymmetric, but similarly-sized halves: 1) a *γ*-TuSC oligomer-like “core” (salmon arrow); and 2) an arrangement of divergent *γ*-TuSC-like subunits (*γ*-TuG4/5 & *γ*-TuG4/6) capped by a terminal *γ*-TuSC that, together, form a large and diverse binding surface for regulatory factors (green arc). H) Helical arrangement of *γ*-tubulin models in the human *γ*-TuRC. I) Helical arrangement of *α*- (white) and *β*- (grey) tubulin dimers within the microtubule lattice.

Lastly, we examined the substantial but unexpected density spanning the *γ*-TuRC lumen, which we named the “lumenal bridge”. This density can be divided into two sections comprising: i) a bi-lobed density located near positions 2 (GCP3) and 3 (GCP2) (Figure 4C); and ii) ∼16 *α*-helices arranged in extended bundles that reach across the lumen and contact positions 9 and 10 (Figure 4C). The bi-lobed density measured ∼5 x 5 x 2 nm and was comprised of a mixture of *α*-helical and *β*-sheet secondary structures. One end of this density contacted the *γ*-tubulins at positions 2 and 3 in the *γ*-TuRC (Figure 4C-D). Remarkably, a model for *β*-actin ^32^ could be rigid-body fitted into this density (Figure 4E), suggesting that this protein adopts an actin-like fold. The presence of an actin-like protein in purified metazoan *γ*-TuRCs has previously been reported ^5,15^, but its potential role in the complex was not known. Our data reveal that an actin-like protein is a structural component of the human *γ*-TuRC, which further exemplifies the diverse structural role for actin-like proteins in large macromolecular assemblies, as in, e.g., the dynactin complex ^33^.

The second section of the lumenal bridge measured ∼9 x 3 x 2 nm and was built from several bundles of consecutive, short (10-20 residues) *α*-helices loosely organized in a chain (Figure S9A). Interestingly, an *α*-helix on one end of this domain contacted the actin-like protein at the barbed end groove (Figure S9A-B), a common interaction site for diverse actin-binding proteins ^34^. The other end of the *α*-helical bundle domain was in close contact with GCP4 at position 9 and GCP5 at position 10 (Figure 4C & F). An unassigned density for a loop stemming from the N-domain of GCP5 (described above) contacted several *α*-helices in these bundles (Figure 4F). Our mass spectrometry data indicated the presence of the small proteins MZT1 and MZT2A/B (Figure S1D) ^13^. Their propensity to form multiple short *α*-helices (Figure S9C-D) leads us to hypothesize that these proteins may constitute at least part of the *α*-helical bundles in this domain. Together, our results identify a structural motif, the lumenal bridge, that spans from the *γ*-tubulins at positions 2-3 via an actin-like protein in the *γ*-TuRC lumen, through a total of >300 Å to the *γ*-tubulins at positions 9-10 on the opposite end of the cone-shaped structure (Figure 4C).

In this study, we determined the first high-resolution structural model of the human *γ*-TuRC, which we find is built from at least 34 distinct polypeptides organized into an asymmetric cone-shaped structure. Our data leads to structural models for human GCP2, GCP3, GCP5, and GCP6 and provide, at near-atomic resolution, many of the critical contacts needed to form this cone-shaped assembly. Our structure reveals that the *γ*-TuRC is built from two key building blocks: 1) the *γ*-TuSC; and 2) subunits that structurally mimic the *γ*-TuSC (i.e., *γ*-TuG4/5 and *γ*-TuG4/6). Together, four *γ*-TuSCs make up one half of the cone, and the *γ*-TuSC-like subunits *γ*-TuG4/5 and *γ*-TuG4/6, ending with a *γ*-TuSC in an altered conformation at the overlap, constitute the other half of the *γ*-TuRC structure. Evolutionary expansion in the metazoan-specific subunits diversifies the *γ*-TuRC (Figure 4G), introducing large (>100,000 Å^2^; typical protein-protein interface: ∼1,000 Å^2 35^), non-*γ*-TuSC surfaces that may interact with specialized “attachment factors” such as NEDD1 ^36^ or hDgt6 ^37^ (e.g., via the GCP6 “plug” (Figure S6)) to establish microtubule polarity ^38,39^.

The structural asymmetry observed in the native *γ*-TuRC was unexpected and has important functional implications. Current models argue that the perfect microtubule nucleation template should match the well-defined helical parameters of the 13-protofilament, 3-start microtubule ^40^. We find that the native human *γ*-TuRC arranges 13 *γ*-tubulins into a single helical turn resembling, but not perfectly matching, the geometry of *α*/*β*-tubulin dimers in the microtubule lattice (Figure 1A and Figure 4H & I) ^41^. It is possible that long-range conformational changes propagating to the “overlap” region - a significant source of structural “breathing” in our overall density map - may be required to facilitate *γ*-tubulin “ring closure” ^40^. Our data suggest that this could occur through the displacement of the *γ*-TuSC at positions 13-14 (Figure 4A-B), the relative movement of inter-*γ*-tubulin interfaces (Figure S4D-E), or changes in the conformation of GCP5 (Figure S5C-F). Alternatively, a helically-perfect ^7^, or even complete ^42^, ring of 13 tubulin-binding sites may not be a prerequisite for the microtubule-nucleating function of the *γ*-TuRC ^3,43,44^.

We find that the ∼2.2 MDa *γ*-TuRC is not solely composed of repeating GCP2 and GCP3 subunits, as current models propose ^17^. The asymmetric, cone-shaped structure observed here is also distinct from similarly-sized, soluble macromolecular complexes, such as the 60S human ribosome (∼25 nm-wide) ^45^, or viral capsids like CCMV (∼20 nm-diameter) ^46^. How might the unusual *γ*-TuRC conical structure form? The human *γ*-TuSC and *γ*-TuSC-like subunits may have a similar propensity to self-assemble as their budding yeast orthologs ^7^, and protein-protein interactions contributed by the lumenal bridge may help to stabilize such assemblies. However, we also note that *γ*-TuRC subcomponents represent some of the least abundant proteins in cultured human cells (<<500 copies per cell ^47^). We propose that a low concentration of compositionally diverse subunits could provide a self-regulating mechanism for limiting the number of *γ*-TuRC assemblies, and, therefore, microtubule initiation sites ^48^, in human cells. Such microtubule “book-keeping” could be important for controlling microtubule architecture and numbers in, e.g., branched microtubule networks ^9^, or in biomolecular condensates in the centrosome ^49^ and liquid-like spindle domains ^8^, which can concentrate tubulin and associated factors to incredibly high levels that would be difficult to regulate ^50^. Our study provides a foundation for dissecting how the *γ*-TuRC assembles and functions in diverse cellular contexts.

## Acknowledgments

The authors are grateful to Dr. G. Alushin for critical reading of the manuscript, as well as M. Ebrahim, J. Sotiris, and H. Ng for fantastic cryo-EM support.

## Methods

### Purification of native human *γ*-TuRCs

Native *γ*-TuRCs were isolated from HeLa S3 cells based on the reported affinity of human *γ*-TuRCs for CDK5RAP2 ^15^.

### Mass spectrometry

A 40 µL aliquot of purified native *γ*-TuRCs was thawed and mixed with 10 µL of 5X sample buffer (Tris-HCl, pH 6.8, 10% (w/v) SDS, 50% (w/v) glycerol, 700 mM 2-mercaptoethanol, and 0.25% (w/v) bromophenol blue). After boiling, the sample was loaded into a single lane of a 4-20% Tris-glycine pre-cast gel with “wide wells” (Novex) and allowed to migrate ∼1 cm into the stacking gel. A corresponding ∼1 cm x 1 cm gel plug was cut out of the gel, further cut into 1 mm cubes, and destained with 33% (v/v) acetonitrile in 50 mM ammonium bicarbonate. To reduce disulfide bonds, gel pieces were incubated with 200 µL 20 mM TCEP in 50 mM ammonium bicarbonate at 56 °C for 45 minutes. Following removal of the TCEP solution, cysteines were alkylated with 200 µL 55 mM 2-chloroacetamide in 50 mM ammonium bicarbonate at room temperature in the dark for 30 minutes, then washed briefly with 50 mM ammonium bicarbonate. Proteins were digested with 80 µL 3.1 ng/uL trypsin in 50 mM ammonium bicarbonate for 4 h at 37°C, followed by addition of a further 125 ng trypsin and incubation overnight. After removing the solution to a clean tube, peptides were extracted from the gel with 100 µL 70% acetonitrile, 1.7% formic acid, 0.1% TFA at 37 °C for 1 h. The supernatant was collected and combined with the initial pool of tryptic peptides. Extraction was repeated sequentially with 100 mM triethyl ammonium bicarbonate and acetonitrile, each with 100 µL at 37 °C for 1 h. Peptides were dried by vacuum centrifugation, resuspended in 5% formic acid, 0.1% TFA, and desalted on C18 membranes before being loaded onto an EASY-Spray column (Thermo Fisher Scientific, 15cm x 75µm ID, PepMap C18, 3µm) via an EASY-nLC 1200 (Thermo Fisher Scientific). Column temperature was set to 35°C. Using a flow rate of 300 nL/min, peptides were eluted with a gradient of 2-32% solvent B in 33 minutes, followed by 32-80% B in 5 minutes, where solvent B was 0.1% formic acid in 95% acetonitrile and solvent A was 0.1% formic acid in water. A spray voltage of 1.8 kV ionized peptides as they eluted into an Orbitrap Fusion Lumos mass spectrometer (Thermo Fisher Scientific) acquiring online data-dependent CID fragmentation spectra.

### Cryo-EM of native *γ*-TuRCs

*γ*-TuRCs were applied to plasma treated Quantifoil R2/2 300-square-mesh copper grids coated with a continuous carbon film, rinsed with washing buffer (40 mM HEPES-KOH pH 7.5, 150 mM KCl, 1 mM MgCl_2_, 0.01% Tween-20 (v/v), 0.1 mM GTP, and 1 mM 2-mercaptoethanol), then blotted and plunge-frozen in liquid ethane using a Vitroblot IV (FEI). Micrographs were recorded on an FEI Titan Krios equipped with a Gatan K2 Summit detector using Serial EM automated data collection ^54^. Data was collected in 3 separate sessions (Figure S2E). The first dataset was exploratory and employed the following settings: 300kV, 70 µm C2 aperture, 33 frames, 45 e^-^/Å^2^, 6.6 s exposure, and 1.335 Å per pixel. In order to acquire as much data as possible, data acquisition was optimized for the subsequent two sessions and the following settings were used: 300kV, 100 µm C2 aperture, 21 frames, 45 e^-^/Å^2^, 4.1s exposure, 1.036 Å per pixel. In these sessions, 8 non-overlapping micrographs were recorded per hole, and a 3 x 3 grid of 9 holes was collected for every stage shift by applying an appropriate beam tilt (i.e., 72 images were recorded per stage shift). This enabled the acquisition of ∼4,500 micrographs per 24 hours in the second and third datasets.

Images for each dataset were corrected for beam-induced drift using MotionCor2 ^55^. CTF parameters for drift corrected micrographs were estimated using CTFFIND4 ^56^. Subsequent processing used Relion v.3.0 ^57^. Initial particle cleanup was conducted using dose-weighted and drift corrected micrographs binned to a pixel size of 5.18 Å (Figure S3). For the first dataset, a small set of particles were manually picked and subjected to reference-free 2D classification. The obtained 2D-class averages that represented particles of different size and shape were used as references for autopicking all binned micrographs from dataset 1. The autopicked particles were cleaned by several cycles of 2D classification. These cleaned particles were used to generate an initial model. The initial model was low-pass filtered to 200 Å and used as a reference for 3D refinement followed by 3D classification. 99,524 particles corresponding to the major 3D class were selected. These particles were subjected to another round of 2D classification in order to select balanced number of particles of different orientations: 35k particles corresponding to the dominant 2D class averages and 35k that corresponded to the rarer views. The obtained 2D class averages were used as references for autopicking from datasets 2 and 3.

As a large number of micrographs were recorded for datasets 2 and 3 (Figure S2E), the autopicked particles were randomly split into subsets of ∼230k particles. In order to avoid losing particles of less abundant orientations, each 230k particle subset was merged with a set of 70k cleaned and balanced particles from dataset 1 (see above). This yielded well resolved 2D class averages during each cycle that pooled particles from the new dataset that might otherwise be lost during classification. After 2 cycles of 2D classification and cleanup, and after removing the 70k particles from dataset 1, the remaining particles (typically ∼100k - 150k) were subjected to 3D refinement. The output translational information from the 3D refinement was used to re-extract particles with the “re-center refined coordinates” option selected in Relion. Re-centered particles were subjected to another round of 3D refinement followed by 3D classification. Particles from 3D classes corresponding to *γ*-TuRC were selected and yielded a set of “good” particles (Figure S2E).

“Good” particles from each dataset were re-extracted as before from the micrographs binned to a pixel size of 1.335 Å. The particles from dataset 1 and 2 were pooled and subjected to global 3D refinement. Due to “breathing” of the complex (Figure S2C), the particles were subjected to another round of local 3D refinement but with a mask including only the stalk densities (excluding globular densities on the open face of the cone). This resulted in better alignment accuracies and, subsequently, higher resolution which in turn yielded better CTF refinement and Bayesian polishing. The CTF-refined and Bayesian-polished particles of datasets 1 and 2 were pooled with “good” particles from the latest dataset (dataset 3) and subjected to another round of 3D classification. A final 467,803 particles were selected from the 3D classes of highest resolution. These “final” particles were used for the overall refinement that yielded 3.81 Å resolution. In order to improve local resolution, the “final” particles were subjected to 3D focused classification and refinement as described before ^58^ (Figure S3).

### Model building

*γ*-tubulin (PDB ID: 3CB2 ^20^) was rigid-body fitted into the overall density map (Figure 1A) using UCSF Chimera’s “Fit in map” function ^59^. *γ*-tubulin models in positions 1-13 were refined into the corresponding density maps using the “Real space refine zone” feature in COOT with torsion, planar peptide, trans peptide and Ramachandran restraints enforced and using a refinement weighting of ≤10 ^60^. GDP was similarly modeled into the *γ*-tubulin nucleotide binding site density.

Models for GCP4 were built by dividing the crystal structure for GCP4 (PDB ID: 3RIP ^22^) into N- (a.a. 1 - 346) and C- (a.a. 347 - 657) domains, which were independently rigid-body fitted into stalk-like density maps at positions 9 and 11. Models were refined into the density maps as described above for *γ*-tubulin. N- and C-domain models were then joined together and the connecting residues were real-space refined in COOT.

Sequences corresponding to the N- and C-domains of GCP2, GCP3, and GCP5 were determined from multiple sequence alignments with GCP4 combined with secondary structure predictions using JPred ^61^. Initial models for GCP2, GCP3, and GCP5 N- and C-domains were generated with the Phyre2 server ^62^ using the GCP4 crystal structure ^22^ as a template. Models were fitted, corrected where necessary using secondary structure predictions and *α*-helical registries of the densities as a guide, and refined into the *γ*-TuRC density as described above for *γ*-tubulin and GCP4.

For GCP6, high quality density in the N-domain was used to identify GCP6 at this position of the complex and to build a corresponding *de novo* N-domain model as described above. For the C-domain, a homology model was generated (Phyre2) with the whole sequence as an input using GCP4 as a template and partially built it into the C-domain density. Then, using secondary structure predictions as a guide, coordinates were *de novo* built and refined (Figure S6A). A homology model for the GCP6 “plug” (a.a. 1-389) was generated based on a top-scoring match to a spliceosomal protein (SF3b155; PDB ID: 5IFE ^28^) from a sequence-profile alignment search using Neff-MUSTER ^27^. The model was rigid-body fitted into the “plug” density with its C-terminus oriented close to the N-terminus of the GCP6 N-domain.

A model for the *γ*-TuNA was built *de novo* in Coot and refined as above. Models for poly-alanine *α*-helices in the lumenal bridge, as well as the CC and HB proximal to GCP6, were built using the “Place helix here” feature in COOT followed by real-space refinement to match the helical twist, as necessary.

### Data availability

The data that support the findings of this study are available from the corresponding author upon reasonable request.

