## Supplementary Information for "Asymmetric molecular architecture of the human *γ*-tubulin ring complex"

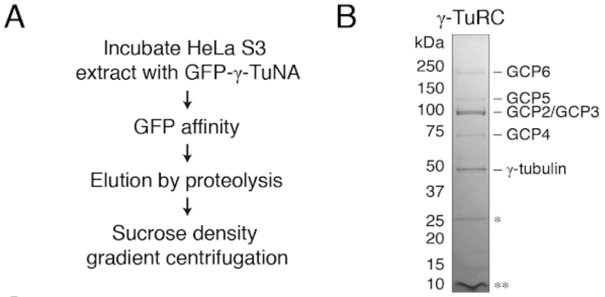


C

Mass spectrometry analysis of γ-TuRC "core" components

| Component | Accession | Peptides (unique/total) | Coverage | iBAQ ratio rel. to GCP6 | Structural stochiometry | Structural identification? |
|---|---|---|---|---|---|---|
| γ-tubulin | P23258 | 10/32 | 83.8 % | 13.8 | 14 | Yes |
| GCP2 | Q9BSJ2 | 61/61 | 72.4 % | 5.3 | 5 | Yes |
| GCP3 | Q96CW5 | 53/53 | 60.4 % | 5.4 | 5 | Yes |
| GCP4 | Q9UGJ1 | 23/23 | 49.1 % | 1.9 | 2 | Yes |
| GCP5 | Q96RT8 | 52/52 | 56.8 % | 0.8 | 1 | Yes |
| GCP6 | Q69RT7 | 51/51 | 40.6 % | 1 | 1 | Yes |

D

Mass spectrometry analysis of γ-TuRC-associated factors

| Component | Accession | Peptides (unique/total) | Coverage | iBAQ ratio rel. to GCP6 | Structural stochiometry | Structural identification? |
|---|---|---|---|---|---|---|
| γ-TuNA | Q96SN8 | 4/4 | 54.0% | 1.6 | 5 | Yes |
| β-actin-like | P60709 | 8/15 | 52.3% | 2.8 | 1 | Yes |
| MZT1 | Q08AG7 | 2/2 | 39.0% | 0.1 | N/A | No |
| MZT2A/B | Q6NZ67 | 2/2 | 27.2% | 0.1 | N/A | No |

**Figure S1. Purification and characterization of native human $\gamma$-TuRC, related to Figure 1.** A) Native human $\gamma$-TuRC purification scheme. B) SDS-PAGE analysis of purified native human $\gamma$-TuRC. Bands corresponding to core $\gamma$-TuRC components are indicated. Bands for GCP2 and GCP3 are very close to each other due to similar molecular weights. Contaminating bands were observed at ~25 kDa and ~10 kDa, likely corresponding to uncleaved GFP-$\gamma$-TuNA (*) and cleaved $\gamma$-TuNA (**) based on size. Double asterisk indicates a contaminant at <10 kDa that likely corresponds to cleaved $\gamma$-TuNA. C) Mass spectrometry analysis of native human $\gamma$-TuRCs showing results for "core" components. iBAQ (intensity-based absolute quantification [16]) ratio was normalized relative to a single copy of GCP6. D) The same mass spectrometry dataset in C) showing results for $\gamma$-TuRC associated factors.

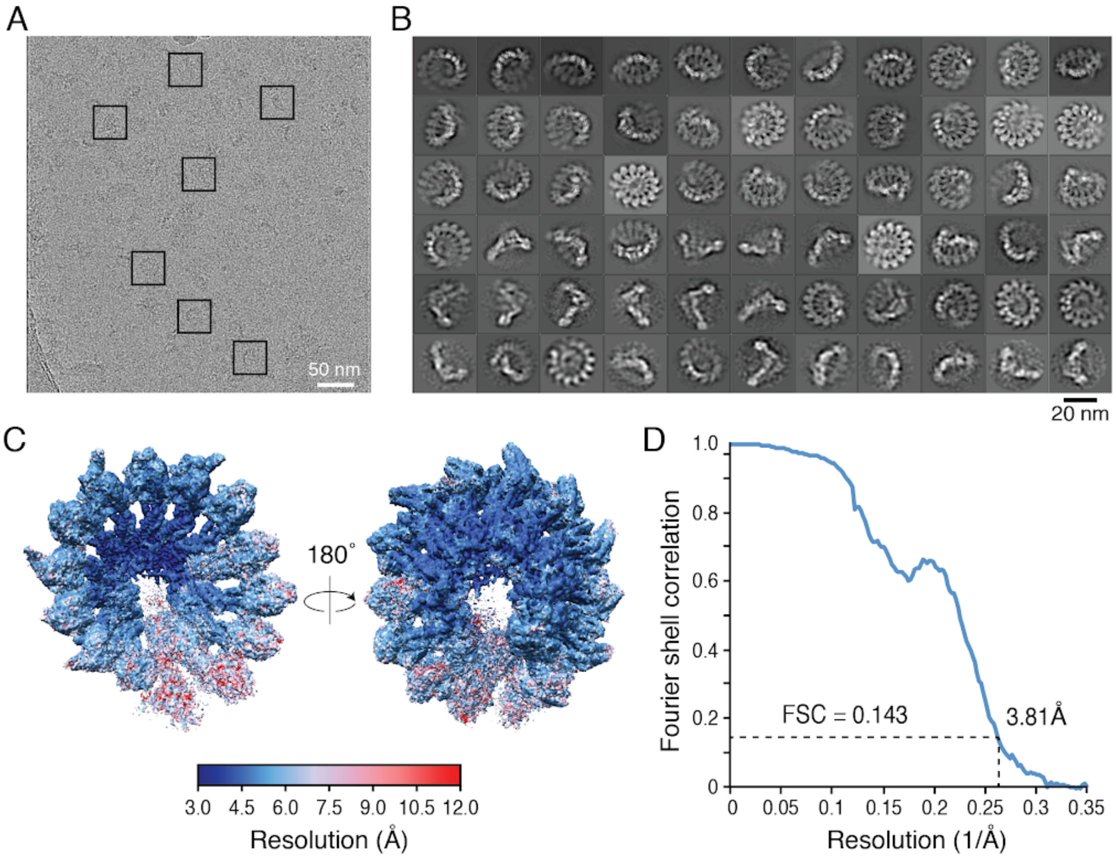


E Data collection statistics:

| | Dataset 1 | Dataset 2 | Dataset 3 | Total |
|---|---|---|---|---|
| Voltage (kV) | 300 | 300 | 300 | |
| Pixel size (Å/pixel) | 1.335 | 1.036 | 1.036 | |
| Exposure time (s) | 9.9 | 4.2 | 4.2 | |
| Total dose ($e^{-}/Å^{2}$) | 45 | 45 | 45 | |
| Number of frames | 33 | 21 | 21 | |
| Micrographs | 4,783 | 42,408 | 22,658 | 69,849 |
| Autopicked particles | 836,778 | 5,182,081 | 2,967,555 | 8,986,414 |
| Good particles | 99,524 | 614,608 | 381,296 | 1,095,428 |
| Final particles | 60,173 | 244,186 | 162,814 | 467,803 |

**Figure S2. Cryo-EM reconstruction of the native human $\gamma$-TuRC, related to Figure 1.** A) Cryo-EM micrograph of native human $\gamma$-TuRCs. Example particles selected for further processing are indicated (boxes). B) 2D-class averages of the $\gamma$-TuRC. C) Two views of the overall $\gamma$-TuRC density map analyzed by ResMap [63], showing a resolution distribution ranging from 3 to 12 Å. D) Gold-standard Fourier shell correlation (FSC) curve of the overall $\gamma$-TuRC density map. The FSC at 0.143 is indicated. E) Cryo-EM data collection and processing statistics used for the refinement of the $\gamma$-TuRC.

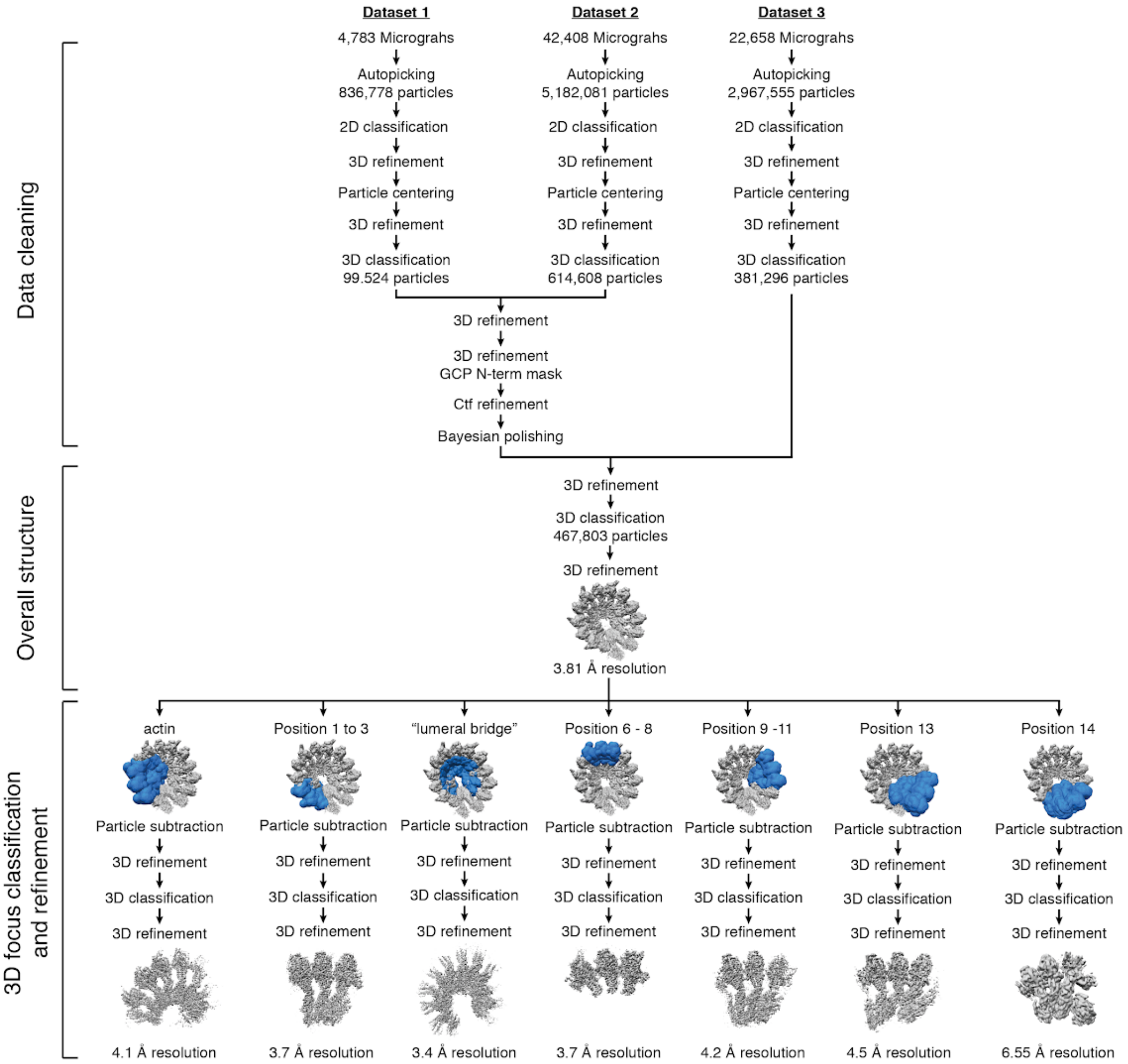


**Figure S3. Cryo-EM data processing of the $\gamma$-TuRC, related to Figure 1-4.** Focused 3D classification and refinement procedures used to improve the resolution in the overall $\gamma$-TuRC density map.

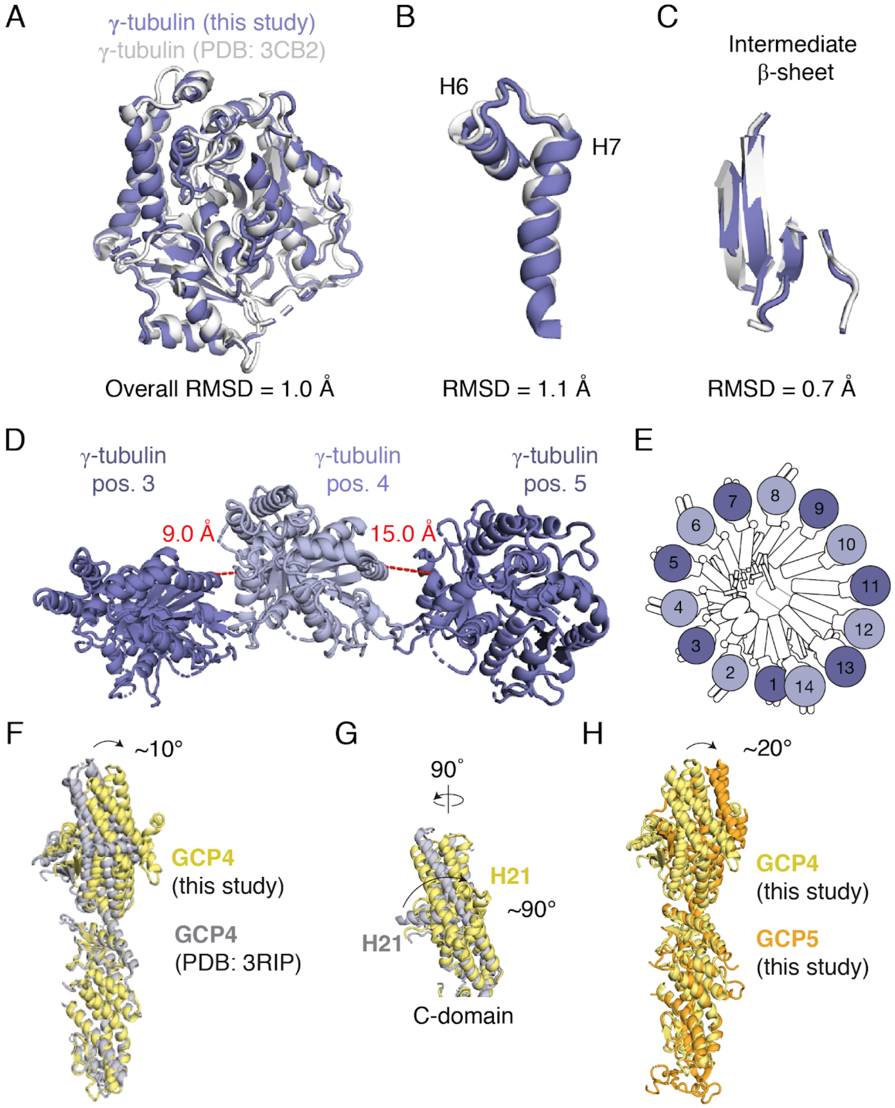


**Figure S4. Comparison of $\gamma$-tubulin and GCP4 to known structures and alignment of GCP5 to GCP4, related to Figure 1 and 2.** A) - C) Alignment of the $\gamma$-tubulin model from this study (blue cartoon representation) to a previously-reported crystal structure of $\gamma$-tubulin:GDP ([20]; white cartoon representation). Alignment was performed as in [20]. RMSD of the overall alignment is indicated in A). B) and C) RMSDs of hallmark $\gamma$-tubulin secondary structure features are indicated. D) View of $\gamma$-tubulin models at position 3-5 highlighting a displacement (red dashed lines, distance indicated) in the $\gamma$-tubulin:$\gamma$-tubulin interface at position 4-5 (light blue - dark blue) compared to at position 3-4 (dark blue - light blue). E) Schematic of the $\gamma$-TuRC highlighting locations of the two types of $\gamma$-tubulin:$\gamma$-tubulin interfaces shown in using the same $\gamma$-tubulin coloring scheme in D). F) Alignment of GCP4 model from this study (yellow cartoon representation) with a previously reported crystal structure (PDB ID: 3RIP; [22]; grey cartoon representation) via their N-domains. A ~10° rotation in the C-domain of our GCP4 model relative to the aligned N-domains is indicated (arrow). G) Rotated view of F) focusing on the C-domain. A ~90° rotation in helix 21 relative to the C-domain is indicated. H) Alignment of our GCP5 model (orange cartoon representation) with our GCP4 model (yellow cartoon representation) via their N-domains. A ~20° rotation in the GCP5 C-domain relative to the aligned N-domains indicated (arrow).

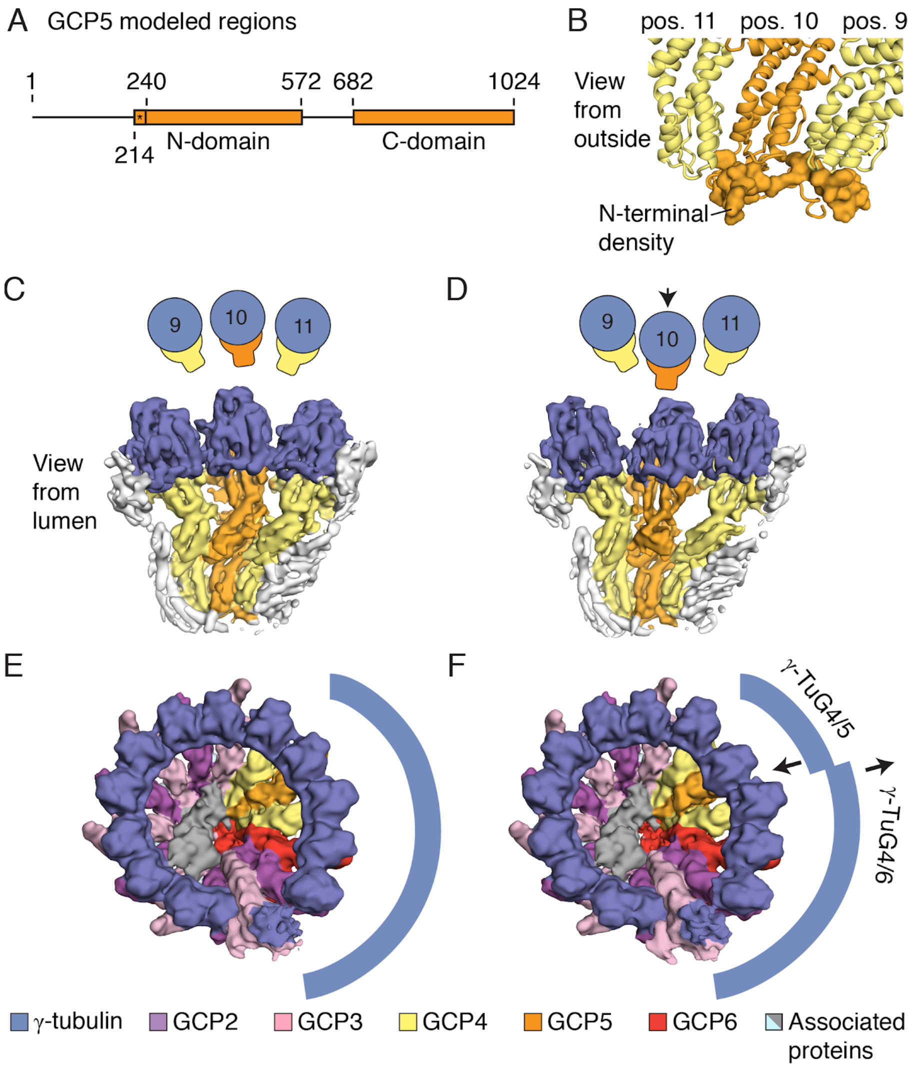


**Figure S5. GCP5 C-domain adopts extended and compacted conformations, related to Figure 2.** A) Schematic of the human GCP5 sequence highlighting regions modeled in this study (orange rectangles). Asterisk refers to GCP5 residues 214-240 modeled with alanine residues only. B) View of the GCP5-associated N-terminal density (orange surface representation) that interacts with N-domains of GCP4 (position 9), GCP5 (position 10) and possibly GCP4 at position 11 (cartoon representations). C) and D) Lumenal view of positions 9-11 of the $\gamma$-TuRC with the GCP5 C-domain in the extended (C) and compacted (D) conformation. Downward displacement in $\gamma$-tubulin at position 10 is indicated (arrow). E) and F) Compaction (F) of the GCP5 C-domain at position 10 results in a displacement of the $\gamma$-TuG4/6 subcomplex away from the conical axis relative to the “elongated” GCP5 subclass shown in (E), represented by the broken arc with arrows showing displacement directions. Maps were low-pass filtered to 8 Å (C-D) and 13 Å (E-F). Unless otherwise indicated, subunits are labeled according to the legend at the bottom of the figure.

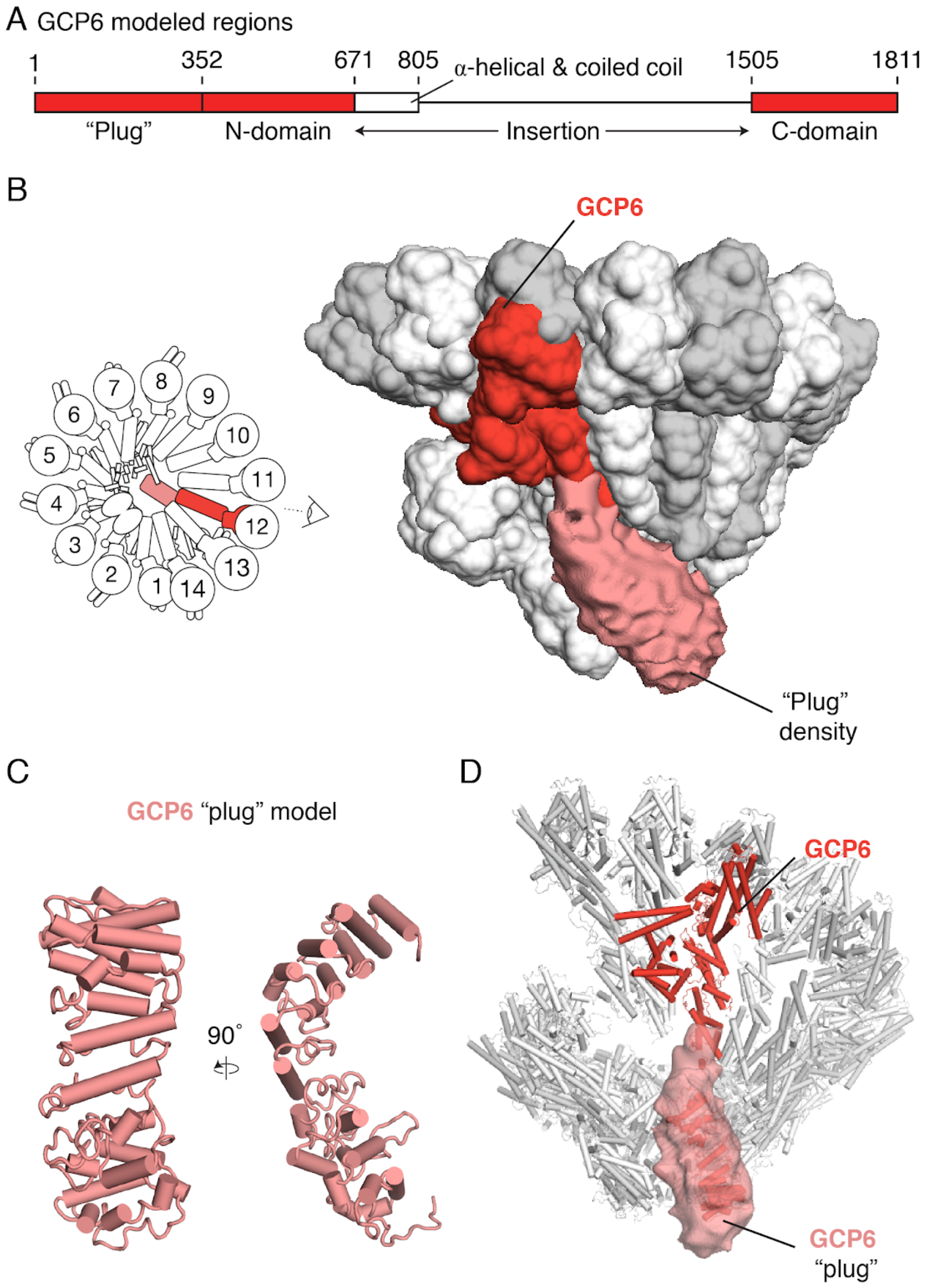


**Figure S6. Model for the GCP6 "plug" related to Figure 2.** A) Schematic of the human GCP5 sequence highlighting regions modeled in this study (red rectangles). The ~800 residue insertion sequence is indicated, including a highly $\alpha$-helical region containing coiled coils (see Methods) shown as a white rectangle. B) Left: Schematic of the $\gamma$-TuRC highlighting the location of GCP6 and establishing the viewing angle. Right: $\gamma$-TuRC model (surface representation) with GCP6 highlighted (red) overlayed with a segmented, 13 Å low-pass filtered density map of the "plug" density (light red surface representation, indicated). C) Cartoon representation of a homology model for the GCP6 "plug". D) Rigid-body fit of the model from C) (light red cartoon representation) into the "plug" density (transparent surface). The GCP6 N- and C-domain model (red cartoon representation) is highlighted in the context of the rest of the $\gamma$-TuRC model (white cartoon representation).

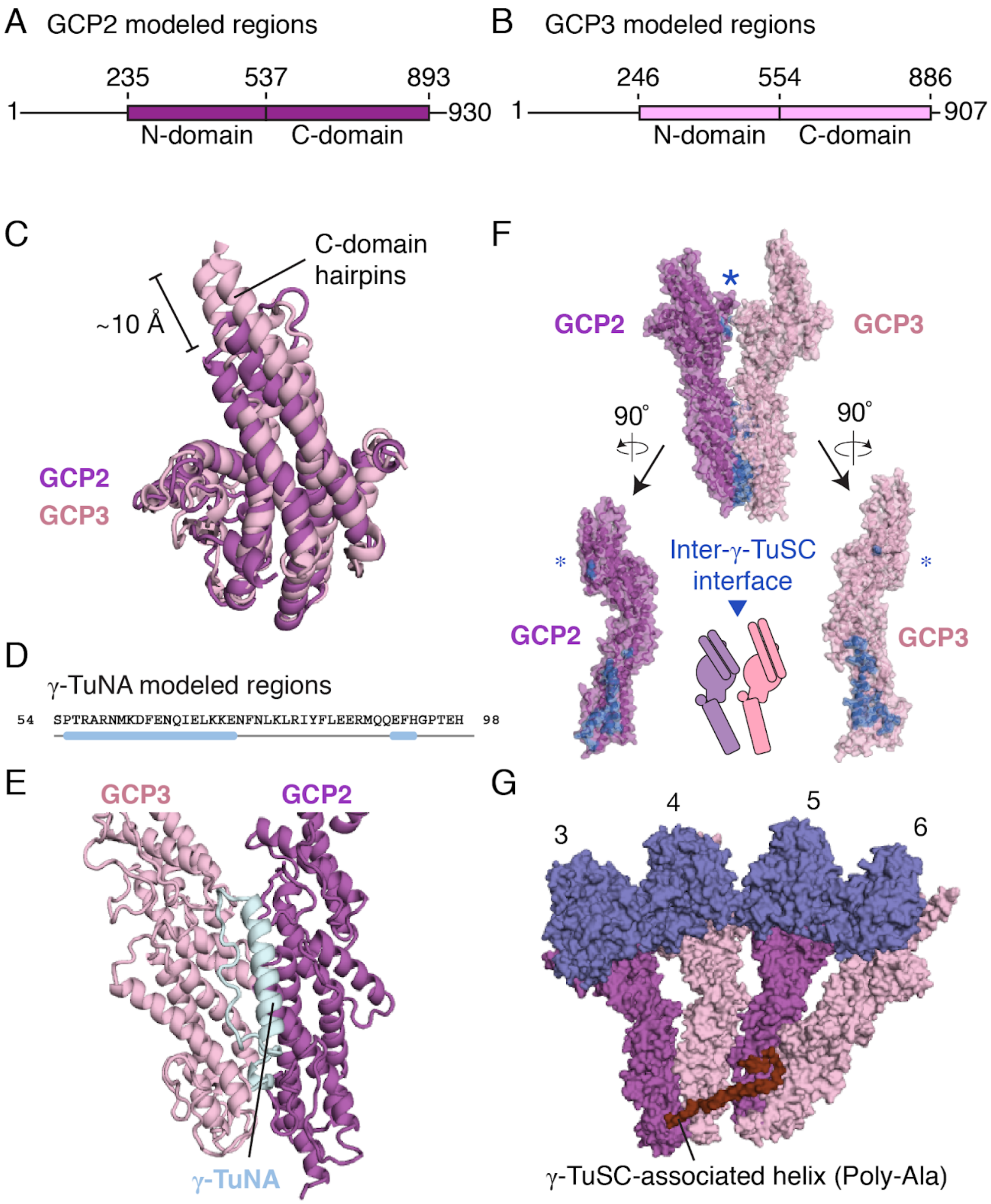


**Figure S7. $\gamma$-TuNA and an extended associated helix stabilize $\gamma$-TuSCs, related to Figure 3.** A) Schematic of the human GCP2 sequence highlighting regions modeled in this study (purple rectangles). B) Schematic of the human GCP3 sequence highlighting regions modeled in this study (pink rectangles). C) Alignment of GCP2 and GCP3 models to one another focusing on the C-domain hairpin. The ~10 Å extension of the GCP3 vs. the GCP2 hairpin is indicated. D) Secondary structure elements of the $\gamma$-TuNA model matched against the primary sequence of the CM1 domain of CDK5RAP2. E) $\gamma$-TuNA is located at the interface between GCP2 and GCP3 N-domains. F) GCP2 (purple surface) and GCP3 (pink surface) models rotated along their long axes (indicated) highlighting the inter-$\gamma$-TuSC interface (blue surface). An asterisk indicates a small interface formed between C-domains that is missing in the intra-$\gamma$-TuSC interface (Figure 3G). G) A long unassigned $\alpha$-helix density (brown surface representation) spans along the N-domains of $\gamma$-TuSCs at positions 3 to 6 (surface representation colored according to Figure 1).

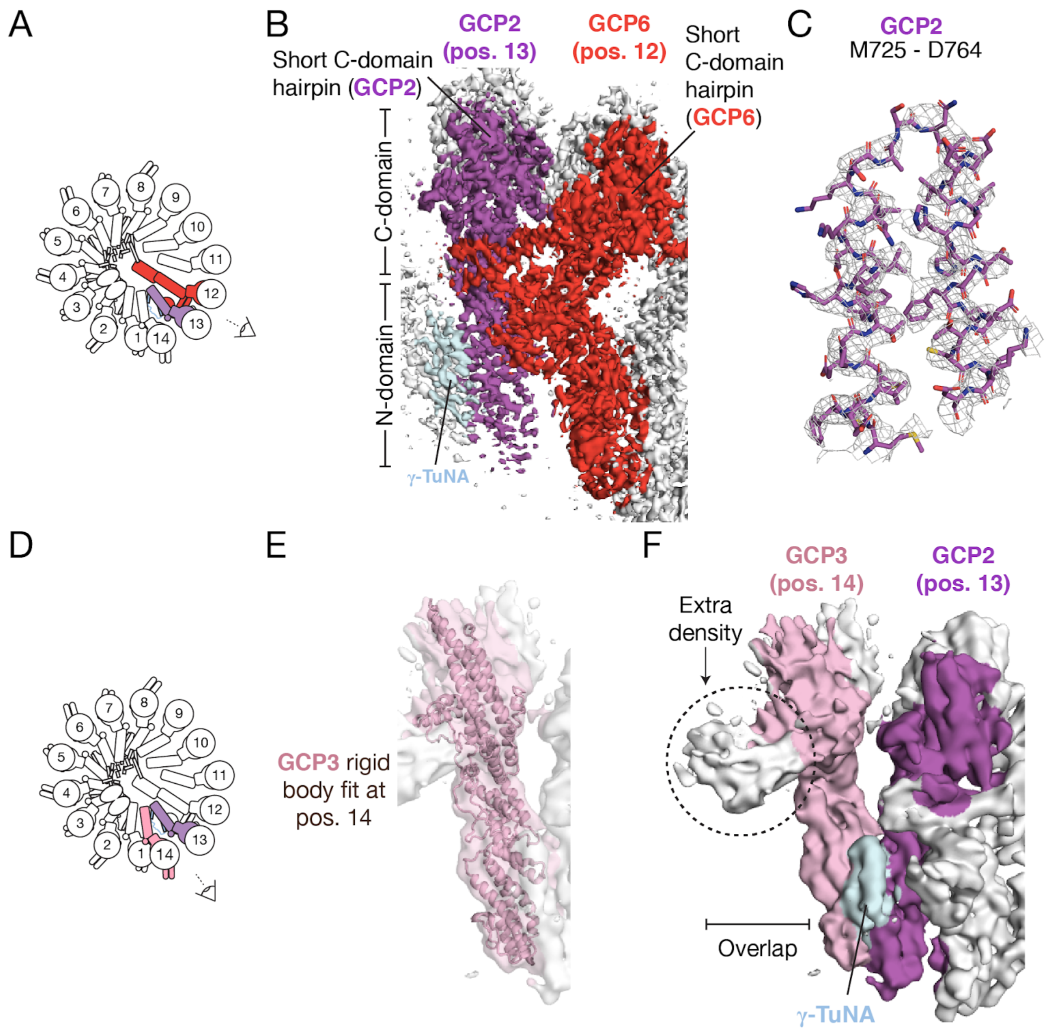


**Figure S8. Identification and organization of a terminal $\gamma$-TuSC at the "overlap" region, related to Figure 4.** A) Schematic of the $\gamma$-TuRC highlighting the locations of GCP6 (red) adjacent to a GCP2 subunit at position 13 (purple) that is associated with a $\gamma$-TuNA (light blue). B) Density map for positions 11-13 used to model GCP2 (position 13, purple surface) with a $\gamma$-TuNA (light blue surface) viewed from the angle indicated in A). C) Relaxed fit of GCP2 C-domain residues M725 - D764 (stick representation) in the corresponding $\alpha$-helical density (mesh) at position 13. D) Schematic of the $\gamma$-TuRC highlighting the location of GCP2 (purple) and a $\gamma$-TuNA (light blue) adjacent to a GCP3 subunit (pink) at position 14. E) Rigid-body fit of the GCP3 model (pink cartoon representation) into the corresponding density (pink transparent surface) at position 14. F) View of GCP2 (purple surface), GCP3 (pink surface), and $\gamma$-TuNA (light blue surface) density constituting the terminal $\gamma$-TuSC at position 13-14. Indicated is an extra density associated with GCP3 (dashed black circle; see also Figure 4A) that is situated above the $\gamma$-tubulin in position 1 (the "overlap"). Views in E - F) are from the angle specified in D).

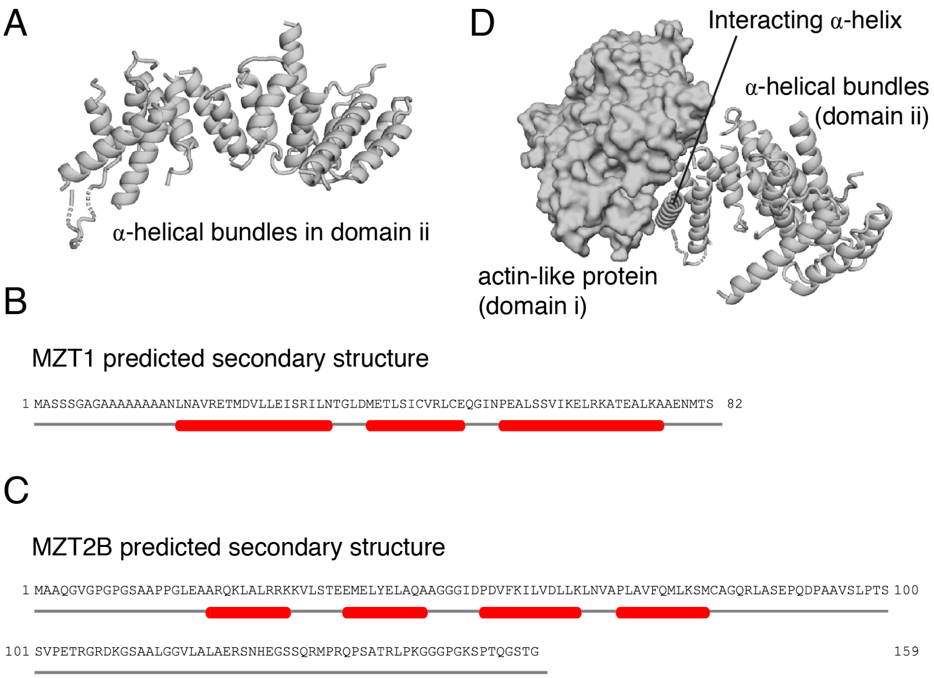


**Figure S9. Features of lumenal bridge domains (i) and (ii), related to Figure 4.** A) A poly-alanine model (cartoon representation) built into the $\alpha$-helical bundle densities comprising domain (ii) of the $\gamma$-TuRC lumenal bridge. An interacting helix that contacts the actin-like protein is indicated. B) $\alpha$-helical bundles of domain (ii) (grey cartoon representation) interacting with a model for the actin-like protein in domain (i) (grey surface representation) in the lumenal bridge. An interacting $\alpha$-helix contacts the actin-like protein at the barbed end groove (indicated). C) and D) Secondary structure predictions of MZT1 and MZT2B ($\alpha$-helical regions shown as red bars).
